# Analysis of Cyp51 protein sequences shows 4 major Cyp51 gene family groups across Fungi

**DOI:** 10.1101/2022.05.18.492509

**Authors:** Brandi N Celia-Sanchez, Brandon Mangum, Marin Brewer, Michelle Momany

## Abstract

Azole drugs target fungal sterol biosynthesis and are used to treat millions of human fungal infections each year. Resistance to azole drugs has emerged in multiple fungal pathogens including *Candida albicans, Cryptococcus neoformans, Histoplasma capsulatum*, and *Aspergillus fumigatus*. The most well-studied resistance mechanism in *A. fumigatus* arises from missense mutations in the coding sequence combined with a tandem repeat in the promoter of *cyp51A*, which encodes a cytochrome P450 enzyme in the fungal sterol biosynthesis pathway. Filamentous members of Ascomycota such as *A. fumigatus* have either one or two of three Cyp51 paralogs (Cyp51A, Cyp51B, and Cyp51C). Most previous research in *A. fumigatus* has focused on Cyp51A due to its role in azole resistance. We used the *A. fumigatus* Cyp51A protein sequence as the query in database searches to identify Cyp51 proteins across Fungi. We found 435 Cyp51 proteins in 301 species spanning from early-diverging fungi (Blastocladiomycota, Chytridiomycota, Zoopagomycota and Mucormycota) to late-diverging fungi (Ascomycota and Basidiomycota). We found these sequences formed 4 major Cyp51 groups: Cyp51, Cyp51A, Cyp51B, and Cyp51C. Surprisingly, we found all filamentous Ascomycota had a Cyp51B paralog, while only 50% had a Cyp51A paralog. We created maximum likelihood trees to investigate the evolution of Cyp51 in Fungi. Our results suggest Cyp51 is present in all fungi with three paralogs emerging in Pezizomycotina, including Cyp51C which appears to have diverged from the progenitor of the Cyp51A and Cyp51B groups.

**Author Summary:** Each year millions of people are infected by a fungal pathogen and receive antifungal treatment with azole drugs. Resistance to azole drugs is becoming increasingly prevalent and is mostly caused by mutations in the azole drug target, Cyp51. *Aspergillus fumigatus* is an airborne fungal pathogen that causes more than 600,000 deaths every year. Azole resistance in *A. fumigatus* is primarily driven by a promoter repeat coupled with mutations in *cyp51A*. In our study, we found 435 Cyp51 proteins in 4 major groups across Fungi, with some species having multiple Cyp51 proteins (Cyp51, Cyp51A, Cyp51B, and Cyp51C). Although most research in *A. fumigatus* has focused on Cyp51A, we found Cyp51B in all filamentous Ascomycota fungi showing it is more conserved than Cyp51A and likely plays a vital role in these fungi.

**Author Summary (Shortened):** Resistance to azole drugs is becoming increasingly prevalent and is mostly caused by mutations in the azole drug target, Cyp51. Azole resistance in Aspergillus fumigatus is primarily driven by a promoter repeat coupled with mutations in cyp51A. We found 435 Cyp51 proteins in 4 major groups across Fungi, with some species having multiple Cyp51 proteins (Cyp51, Cyp51A, Cyp51B, and Cyp51C). Although most research focuses on Cyp51A, we found Cyp51B in all filamentous Ascomycota fungi showing it’s more conserved than Cyp51A.

## Introduction

Fungal pathogens caused over 9 million diagnosed infections in 2017 in the United States, but the true fungal burden is hard to estimate since many cases are likely undiagnosed [1, 2]. The infections caused by fungal pathogens include severe chronic conditions, complex chronic respiratory conditions, recurrent infections, and many life-threatening invasive diseases [3]. Invasive fungal infections generally occur in individuals with suppressed or compromised immune systems [4]. These infections have a high mortality rate if not treated early with appropriate antifungal drugs [4]. Major drugs used to treat invasive fungal infections are echinocandins, polyenes, flucytosine, and azole drugs [5]. Azoles, which target synthesis of the fungal-specific membrane component ergosterol, are among the most highly used antifungal drugs.

Cyp51 proteins, also known as Erg11 in Ascomycota yeast in the Saccharomycotina and Taphrinomycotina subphyla, are in all biological kingdoms and are highly conserved [6]. Cyp51 proteins have 6 substrate recognition sites (SRS), an oxygen-binding motif (AGXDTT), PER and EXXR motifs that create an E-R-R triad within the heme pocket, and a conserved heme-binding motif (FXXGXXXCXG) (Fig. 1) [7, 8]. Azole drugs competitively bind to sterol 14 alpha-demethylase (Cyp51, Erg11), a cytochrome P450 in the ergosterol biosynthesis pathway in fungi. Azole drugs consist of a heterocyclic ring with either two (imidazoles) or three (triazoles) nitrogens and a sidechain. The side chain of azoles interacts with the Cyp51 polypeptide while the nitrogen in the azole heterocyclic ring interacts directly with the sixth ligand of the heme ferric ion, a cofactor of the Cyp51 protein [9]. Cytochrome P450 proteins conduct a three-step reaction within the sterol biosynthesis pathway leading to the production of cholesterol in animals, sitosterol in plants, and ergosterol in fungi [10, 11]. Sterols are integrated into the cell membrane where they aid in membrane fluidity and permeability [10]. The binding of azoles to Cyp51 depletes intracellular ergosterol and causes accumulation of methylated sterols and toxic intermediate sterols within the fungal cell membrane causing arrested growth and cell membrane stress [12].

**Figure 1:**
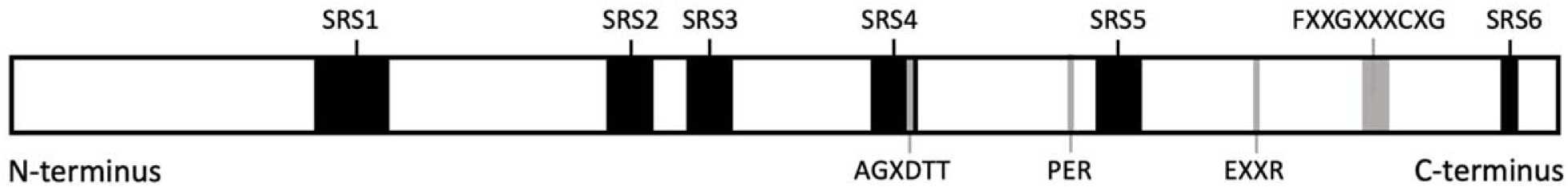
Typical organization of Cyp51 domains. Cyp51 proteins contain six substrate recognition sites (SRS 1-6), an oxygen binding motif (AGXDTT), PER and EXXR motifs, and a conserved heme-binding motif (FXXGXXXCXG). Black boxes represent SRS domains. Gray shading represents other motifs. Diagram is based on *A. fumigatus* Cyp51A (XP_752137.1) and is shown to scale.

Many fungi have acquired mutations in *cyp51*that alter the ability of azoles to bind and inhibit Cyp51 [13]. Many missense mutations that have been shown to decrease sensitivity to azoles as determined by minimum inhibitory concentration (MIC) in the human fungal pathogens *Candida albicans, Cryptococcus neoformans, Histoplasma capsulatum*, and *A. fumigatus* [13-17] occur in substrate recognition sites causing azoles to interact and bind differently within Cyp51. Increased expression levels of *cyp51A* due to 34-, 46-, 53- and 120-bp tandem repeats in the promoter have occurred in *A. fumigatus* leading to high levels of pan-azole resistance, resistance to more than one azole drug [18-21]. Tandem repeats in the *cyp51A* promoter reduce the affinity of the promoter and the CGAAT binding complex (CBC), which binds to CGAAT in the promoter and downregulates *cyp51A* expression [22]. Although Cyp51A has been the focus of most studies in *A. fumigatus*, a second paralog (Cyp51B) has also been documented to cause resistance through upregulation and missense mutations [23, 24]. Like human pathogens, plant pathogens (such as *Erysiphe necator, Mycosphaerella* spp., *Penicillium digitatum, Venturia inaequalis*) undergo changes in the *cyp51A* promoter (substitutions, insertions, duplications) and/or mutations in substrate recognition sites to alter expression and binding of Cyp51A [25-34].

Filamentous Ascomycota have multiple Cyp51 paralogs that may play different roles in the cell under normal conditions or under azole stress. The goal of our study was to understand the evolutionary relationships among fungal Cyp51 proteins.

## Results and Discussion

### 435 fungal Cyp51 proteins were analyzed

Cyp51 proteins were previously defined as having 6 substrate recognition sites (SRS), an oxygen-binding motif (AGXDTT), PER and EXXR motifs that create an E-R-R triad within the heme pocket, and a conserved heme-binding motif (FXXGXXXCXG) [7, 8] (Fig. 1). To understand Cyp51 genes across Fungi, the *A. fumigatus* Cyp51A protein was used as a reference in a protein BLAST (Basic Local Alignment Search Tool) [35]. A total of 4404 protein sequences resulted and were filtered to retain those with greater than 50% coverage and greater than 30% percent identity to the reference sequence, resulting in 480 sequences (Supplemental Table 1). The resulting protein sequences were analyzed for the presence of full length SRS1-6 domains and the four Cyp51 motifs. Of these, 439 proteins had SRS1-6 domains, the oxygen-binding motif AGXDTT, the PER and EXXR motifs, and the conserved heme-binding motif FXXGXXXCXG (Supplemental Table 1) and were considered to be functional Cyp51 proteins. Within the 439 Cyp51 proteins, we found 19 fusion proteins in various Ascomycota in which a kinase immediately upstream and Cyp51B were mistakenly fused in genome processing. According to the Joint Genome Institute (JGI), a frameshift mutation occurred *in silico* causing the stop codon of the kinase to appear to be deleted (https://img.jgi.doe.gov/data-processing.html). The Cyp51B portions of the fusion proteins were extracted based on their Cyp51 domains and kept in the analyses. Four Cyp51B proteins (XP_007688940.1, XP_014555703.1, XP_033384229.1, XP_018700143.1) were not included in the analyses due to missing amino acids in SRS1-6 or missing amino acids in conserved motifs resulting in a total of 435 Cyp51 proteins retained for further analysis.

### Fungal Cyp51 proteins fall into 4 groups

To understand presence of Cyp51 homologs across Fungi, we analyzed 435 Cyp51 proteins (Supplemental Table 2). Most fungal species in our study had one or two Cyp51 paralogs (180/295 and 100/295, respectively) (Table 1). Fewer had three (14/295) and only one fungus, *Basidiobolus meristosporus*, had four copies of Cyp51 (Table 1). Fungi with one or two Cyp51 proteins were found across all taxonomic groups (Table 1). Fungi with three Cyp51 proteins were found in Basidiomycota and Ascomycota. One species in Zoopagomycota had four Cyp51 proteins (Table 1). Members in Pezizomycotina had different combinations of Cyp51 paralogs (Table 2). Most species had Cyp51A and Cyp51B paralogs or only Cyp51B (69/171 and 63/171, respectively) (Table 2). All members of Pezizomycotina contained a Cyp51B paralog (Table 2). As shown in Supplemental Table 2, we named fungal Cyp51 proteins without a designation in NCBI based on the group assignment in Supplemental Figure 1.

**Table 1.**
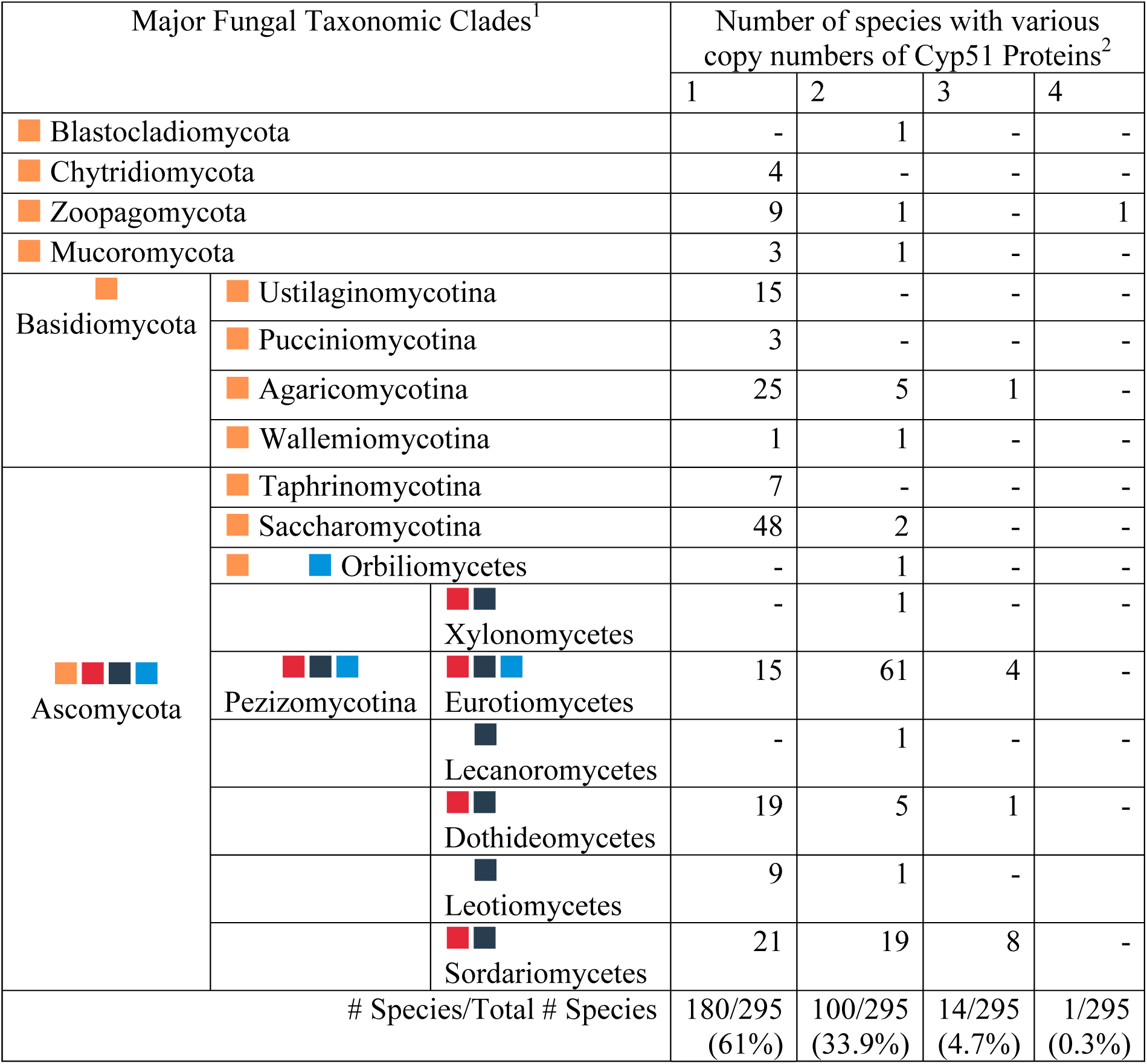
Species of Fungi have different numbers of Cyp51 proteins. Orange, red, black, and blue boxes represent the presence of Cyp51, Cyp51A, Cyp51B, and Cyp51C within fungal groups, respectively. ^1^Major fungal taxonomic clades based on James et al 2020 [36]. ^2^Based on all fungal Cyp51 proteins identified in NCBI databases (435) as described in methods and shown in Supplemental Table 2.

**Table 2.** Species in Pezizomycotina have different combinations of Cyp51 paralogs.

| Cyp51 Paralog | B | A+B | B+B | B+C | A+B+C | A+A+B | A+B+B | B+C+C |
| --- | --- | --- | --- | --- | --- | --- | --- | --- |
| # Species/Total # Species | 63/171 | 69/171 | 4/171 | 18/171 | 9/171 | 5/171 | 1/171 | 1/171 |

To investigate how fungal Cyp51 proteins are related to each other, we used RAxML to create a maximum likelihood tree with 435 fungal Cyp51 proteins and 2 human Cyp51 proteins to root the tree (Supplemental Figure 1). To aid in visualization, we collapsed branches based on phyla, subphyla or classes (Figure 2). We found fungal Cyp51 proteins fell into four major groups which we designated as Cyp51, Cyp51A, Cyp51B, and Cyp51C based on naming in previous literature (Figure 2, Supplemental Figure 1). Cyp51 in members of Saccharomycotina and Taphrinomycotina are also known as “Erg11” in the literature. The topology of our Cyp51 protein tree largely followed the topology of the fungal tree of life (Figure 2, Supplemental Figure 1, [36]). Proteins from early-diverging fungi (Blastocladiomycota, Chytridiomycota, Zoopagomycota and Mucormycota), Basidiomycota, Saccharomycotina, and Taphrinomycotina fell into group Cyp51. Our phylogenetic analyses show a divergence of three Cyp51 paralogs in filamentous Ascomycota (Figure 2, Supplemental Figure 1). Those from Pezizomycotina fell into Cyp51A, Cyp51B, and Cyp51C. Divergence of Cyp51 from the common ancestor of paralogs Cyp51A, Cyp51B and Cyp51C has strong support (100% bootstrap support), but divergence of paralogs Cyp51A, Cyp51B and Cyp51C from each other does not have strong support (41%, 36%, and 41% bootstrap support, respectively).

**Figure 2:**
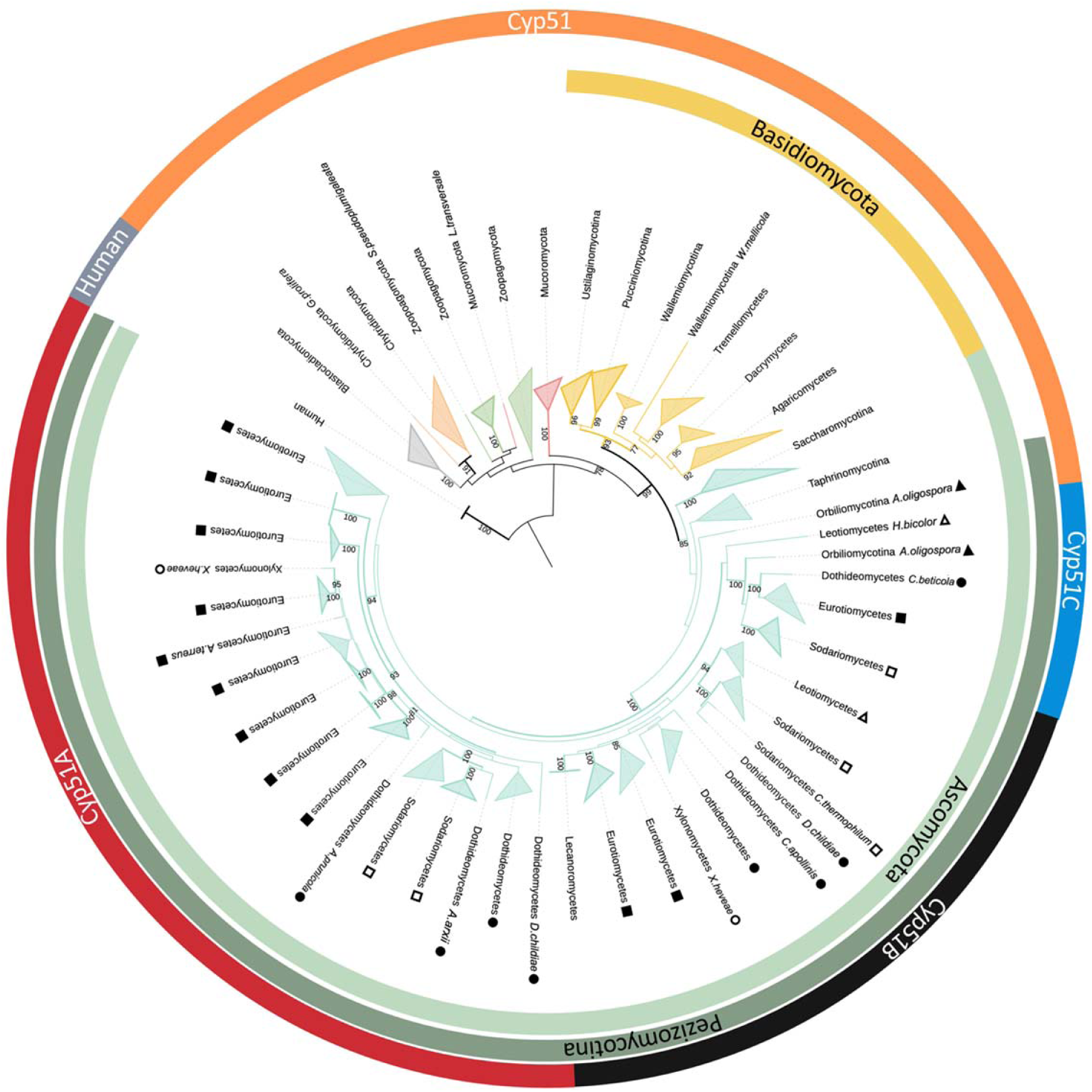
Collapsed Cyp51 protein tree for Fungi. Branch colors match the colors used for taxonomic clades in the Fungal Tree of Life [36]. Branches for Blastocladiomycota, Chytridiomycota, Zoopagomycota, Mucoromycota, Basidiomycota, and Ascomycota are represented by gray, orange, green, red, yellow, and teal, respectively. Branches with bootstrap support of at least 90 are in bold. Collapsed branches represent phyla, subphyla and classes and are named accordingly. Shapes represent subphyla and classes in Ascomycota. Filled triangles represent subphylum Orbiliomycotina. Empty triangles, filled circles, empty circles, filled squares, and empty squares represent classes Leotiomycetes, Dothideomycetes, Xylonomycetes, Eurotiomycetes, and Sodariomycetes, respectively. The most inner ring shows phyla Basidiomycota (yellow) and Ascomycota (teal). The second ring shows subphylum Pezizomycotina (dark teal). The outer ring shows human Cyp51 proteins used as the outgroup (grey) and 4 groups of fungal Cyp51 proteins – Cyp51, Cyp51C, Cyp51B, and Cyp51A represented by orange, blue, black and red, respectively.

We postulate two possible evolutionary paths for Cyp51 paralogs as shown in Figure 3. In the first possible evolutionary path, shown in 3A, after an initial Cyp51 duplication paralog C diverged followed by another duplication event and divergence of paralogs A and B. The divergence of paralogs A (41%), B (36%), and C (41%) has low support. In the second possible evolutionary path, shown in 3B, the poorly supported nodes are removed so that the three paralogs diverged after two unresolved duplication events or a triplication event placing them on the same branch. In either scenario it is possible that subsequent gene loss(es) or duplication(s) led to species with the different combinations of paralogs shown in Table 2.

**Figure 3.**
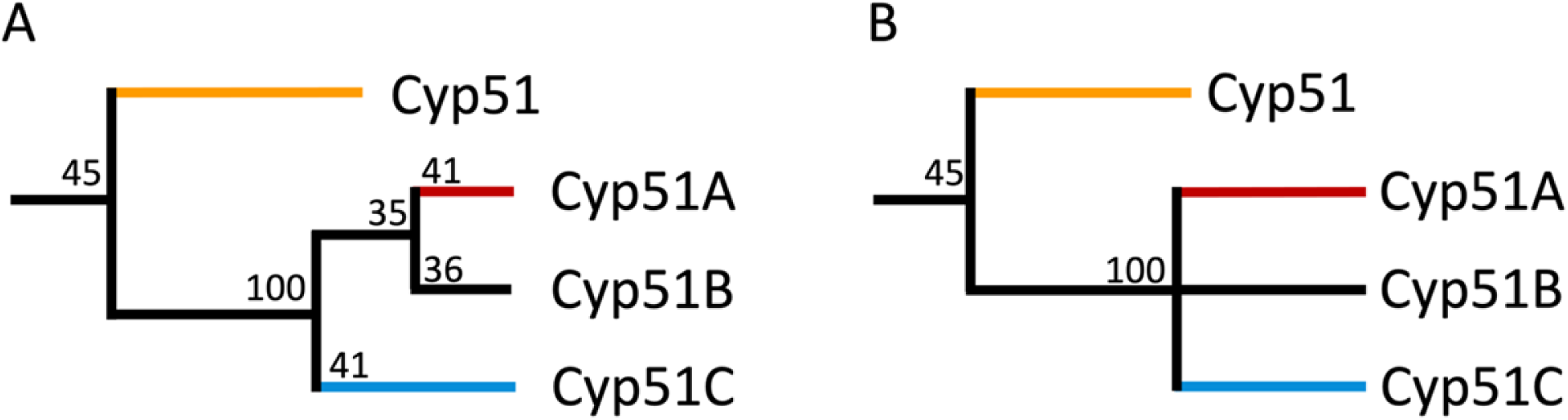
Possible Cyp51 homolog evolutionary paths. Simplified diagrams of possible Cyp51 evolutionary paths based on Figure 2. The Cyp51 branch in early diverging fungi, Basidiomycota, Saccharomycotina and Taphrinomycotina is colored orange. Filamentous Ascomycota (Pezizomycotina) Cyp51A, Cyp51B, and Cyp51C branches are colored red, black and blue, respectively. Numbers represent bootstrap support.

Distinguishing between these possible evolutionary paths is complicated by the small number of Cyp51C sequences and the relatively low number of characters (414-624 amino acids in Cyp51 genes) resulting in low bootstrap support for some nodes. Our analysis only had 29 Cyp51C sequences compared to Cyp51A and B with 87 and 171 sequences, respectively. To see if we could better resolve the relationships among Cyp51A, Cyp51B, and Cyp51C, we analyzed pairwise conservation of all Cyp51 protein sequences using Geneious Prime (Supplemental Table 3). We found that individual members of the Cyp51 group varied the most from each other, with only 46% similarity. Similarity within the Cyp51A, Cyp51B, and Cyp51C groups was much higher (64.7%-68.8%). Comparing between groups, members of the Cyp51 group were 45-50% similar to members of Cyp51A, Cyp51B, or Cyp51C groups while members of Cyp51A, Cyp51B, and Cyp51C groups were roughly 60% similar to each other., (Supplemental Table 3). We then examined conservation within the highly conserved motifs (SRS 1-6, AGXDTT, PER, EXXR and FXXGXXXCXG) (Supplemental Table 4). Once more the general trend was that motifs within the Cyp51 group were more variable that those in other groups. .

Consensus sequences from the four Cyp51 groups were compared to each other to create a visual representation of differences in motifs across groups (Figure 4, Supplemental Table 4). The most conserved regions (>95% across all Cyp51 proteins) are in the EXXR, PER, and FXXGXXXCIG motifs (Figure 4, Supplemental Table 4). These three motifs are presumably highly conserved due to their roles in Cyp51 structure and function. The **E**XX**R** and PE**R** motif form the **E**-**R**-**R** triad that stabilizes the core structure of Cyp51, while the FXXGXXXCIG motif is a heme-binding domain that is essential for Cyp51 function [37, 38]. Out of the 17 amino acids within these three motifs, 13 are more than 95% conserved (Figure 4, Supplemental Table 4). Looking at all motifs, there are 59 amino acids shared between the four consensus sequences with 39 having greater than 95% conservation (Figure 4, Supplemental Table 4). Cyp51A and Cyp51B have the highest amount of shared amino acids in motifs (88%) (Figure 4, Supplemental Table 4). Cyp51C has 23 amino acids across motifs that are unique compared to 15 in Cyp51, 8 in Cyp51A, and 5 in Cyp51B, suggesting Cyp51C may be a specialized group, though the low number of Cyp51C sequences included in the analysis might also explain the pattern (Figure 4).

**Figure 4.**
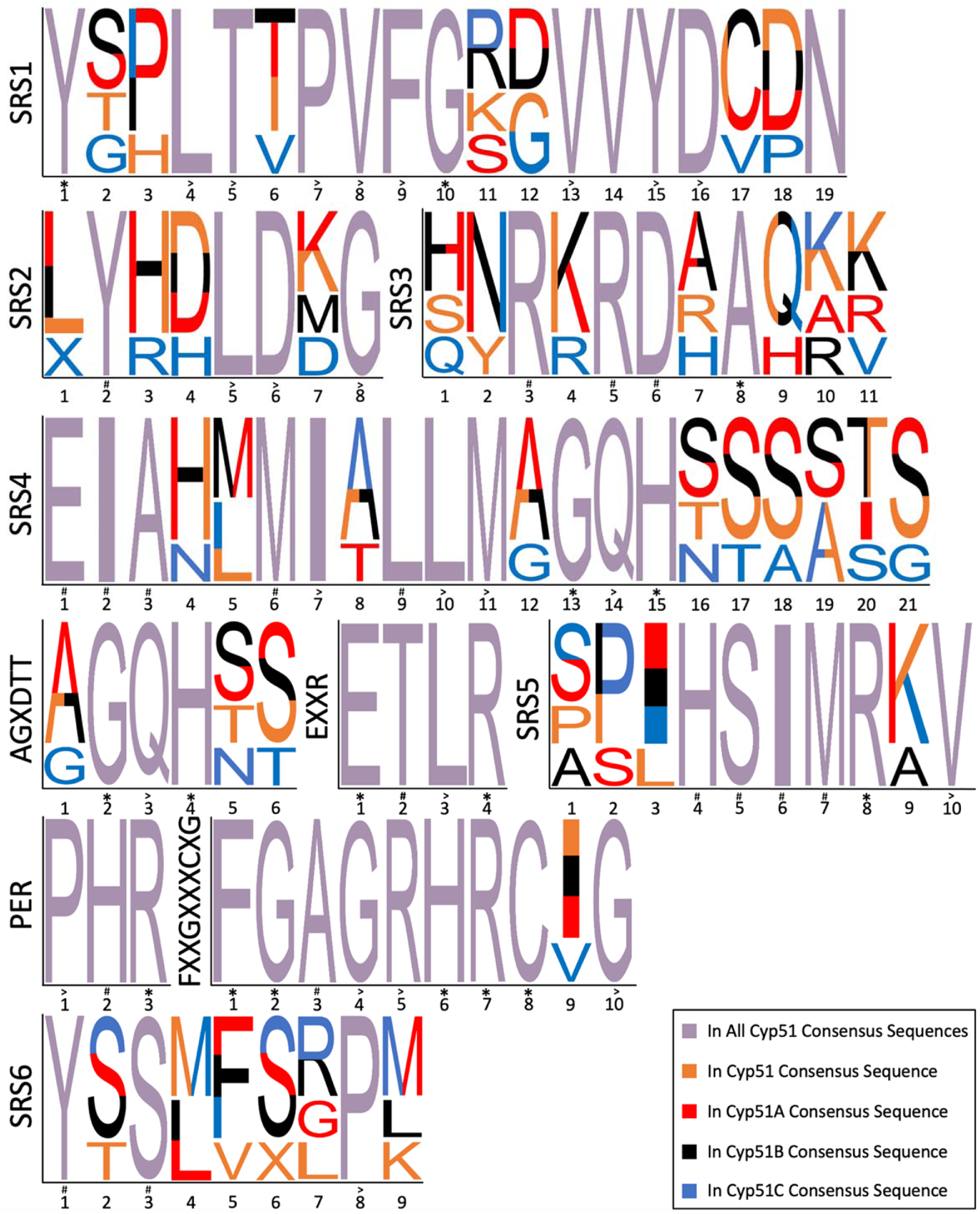
Conservation of motifs in Cyp51 proteins across Fungi. Consensus sequences for each of the four Cyp51A groups are shown arranged from the N- to C-terminus. Amino acids conserved across all 435 Cyp51 protein sequences are denoted by an *. Amino acids found in more than 95% of all Cyp51s are denoted by >. Lower than 95% conservation in all Cyp51s are denoted by #. Numbers represent amino acid position within motifs.

Research on Cyp51 in filamentous Ascomycota tends to be focused on the Cyp51A paralog because of its role in azole resistance in fungal pathogens of plants and animals. To our surprise, we found Cyp51A in only half of the species of filamentous Ascomycota we analyzed (86/171) and Cyp51B in all species of filamentous Ascomycota (Table 2, Supplemental Table 2). The low bootstrap support we saw for divergence of Cyp51A from Cyp51B suggests they may play very similar roles with Cyp51B being the essential paralog and the possible Cyp51 ortholog (Figure 3A). Indeed, in *A. fumigatus*, Cyp51A and Cyp51B act in a compensatory manner; when one paralog is knocked out expression of the other paralog increases [39]. Our results confirm and expand previous work by Hawkins et al [28] that compared 86 fungal Cyp51 proteins in 50 different species and found all species of filamentous Ascomycota retained a Cyp51B paralog, but Cyp51A had been lost in multiple lineages, and Cyp51C was only found in *Fusarium* spp. [40-42]. [40, 41].) . Cyp51C was subsequently reported in *Fusarium spp*., *Gibberella zeae*, and *Nectria haematoccoa* [40, 41]. We found Cyp51C in 9 other genera (Supplemental Table 2). Interestingly, all are pathogens of plants or animals (Supplemental Table 2). Cyp51C has been shown to be necessary for invasion of plant host tissues in *Fusarium* [43]. This raises the interesting possibility that it could play a similar role in other genera, though further functional studies are needed to test the role of Cyp51C in invasion and virulence.

Maximum likelihood analysis suggested two possible evolutionary paths for Cyp51 paralogs:1) Cyp51C diverged before Cyp51A and Cyp51B (Figure 3A; 2) Cyp51A, Cyp51B, and Cyp51C diverged from each other at the same time (Figure 3B). Based on the higher amino acid conservation between Cyp51A and Cyp51B, the evolutionary path shown in Figure 3A seems more likely; that is to say Cyp51C is a specialized group of proteins that diverged first (Figure 3A, Figure 4, Supplemental Table 4). More functional studies are needed to understand shared and unique roles Cyp51A, Cyp51B, and Cyp51C paralogs play in fungi.

## Materials and Methods

### NCBI Protein Blast

Cyp51 protein sequences (XP_006955920.1, XP_016612382.1, XP_025190100.1, XP_01661282.1, XP_018263285.1, XP_752137.1, XP_018249823.1, XP_018226934.1, XP_015469227.1) were used in an NCBI Protein Blast to search the refence sequence database for other Cyp51 proteins in Fungi (Supplemental Table 1). The following setting were used for the Protein BLAST: Database: Reference Proteins, Exclude: uncultured/environmental sample sequences, Algorithm: blastp (protein-protein BLAST), Max Target Sequences: 1000, Expect Threshold: 0.001, Word size: 6, Max matches in a query range: 0, Matrix: BLOSUM62, Gap Costs: Existence: 11 Extension: 1, and Compositional adjustments: Conditional compositional score matrix adjustment. Some clades were not represented in the reference sequence database, so identical settings were used to search the non-redundant proteins sequences database. The unfiltered searches resulted in a total of 4404 sequence hits. Sequences with less than 50% coverage and less than 30% percent identity were eliminated. FASTA files of the 480 sequences resulting from filtering were downloaded and opened in Geneious Prime 2019.1.1. To confirm the sequences were a Cyp51, sequences were checked for the presence of SRS1-6, the oxygen-binding motif AGXDTT, PER and EXXR motifs, and the conserved FXXGXXXCXG heme-binding domain.

The following sequences were eliminated for missing amino acids in SRS1-6 or missing amino acids in conserved motifs: EPZ30787.1, EPZ31936.1, KXN68292.1, RKP07181.1, RKP16598.1, RKP17874.1, RKP18653.1, RKP18926.1, XP_001218650.1, XP_002563403.1, XP_002583031.1, XP_002842283.1, XP_003005233.1, XP_003325369.2, XP_007375289.1, XP_007756389.1, XP_007802603.1, XP_008039623.1, XP_009649122.1, XP_013258864.1, XP_015404015.1, XP_018230821.1, XP_018249826.1, XP_018270027.1, XP_018712692.1, XP_020066776.1, XP_022578172.1, XP_025599710.1, XP_027619241.1, XP_027619242.1, XP_031034290.1, XP_031059536.1, ORZ32486.1, XP_017991977.1, XP_003017020.1, XP_003019064.1, and XP_033461214.1. The following 19 fusion proteins were found in various members in Ascomycota: XP_022511803.1, XP_013278994.1, XP_024670717.1, XP_031899359.1, XP_031927262.1, XP_026621438.1, XP_025554838.1, XP_018192447.1, XP_031935516.1, XP_025433272.1, XP_015404994.1, XP_024709601.1, XP_007688940.1, XP_014073145.1, XP_025394250.1, XP_014555703.1, XP_007712864.1, XP_033384229.1, XP_018700143.1. Four fusions (XP_007688940.1, XP_014555703.1, XP_033384229.1, XP_018700143.1) were not included in the analyses due to missing amino acids in SRS1-6 or missing amino acids in conserved motifs. *Aspergillus flavus* contained three Cyp51 proteins, but were later identified to be contaminated with foreign sequence by NCBI (https://www.ncbi.nlm.nih.gov/protein/XP_002375123.1/) and were removed (XP_002375123.1, XP_002379130.1, XP_002383931.1).

### Phylogenetic analyses

Protein sequences were aligned once with MAFFT version 7.407 then once with PASTA version 1.8.5 [44, 45]. Maximum likelihood trees were constructed with RAxML version 8.2.11 with a PROTGAMMAAUTO or GTRGAMMA substitution model and 1000 bootstraps [46]. Interactive Tree of Life (iTOL) was used for visualization and annotation of the trees [47].

### Amino Acid Analyses

Geneious Prime (version 2021.2.2) was used to generate pairwise identities and consensus sequences. Similarity tables for the whole protein and motifs are based on pairwise identity in each group, number of similar amino acids divided by the total number of amino acids in the protein or motif. “Weblogo-like” diagrams were created manually for visualization of conservation across groups. The height of one letter amino acid designation was based on frequency across all four consensus sequences. Colors and symbols were used as described in Figure 4 legend to denote conservation within groups.

## Data availability

All Cyp51 sequences used are listed in Supplementary Table 2 and are publicly available through NCBI (https://www.ncbi.nlm.nih.gov/).

